# Beliefs and attitudes towards child epilepsy: A structural equation model

**DOI:** 10.1101/2020.01.30.927681

**Authors:** Luanna G. Silva, Izabel C. S. L. Beltrão, Paulo F. R. Bandeira, Gyllyandeson A. Delmondes, Cicero D. Alencar, Simone S. Damasceno, Marta R. Kerntopf

## Abstract

Infantile epilepsy is a chronic neurological disease permeated by negative beliefs and attitudes that strengthen social stigma and discrimination, as well as profoundly limit the routine of the affected child. Therefore, the objective of this study was to analyze the possible influencing factors of the beliefs and attitudes towards childhood epilepsy among users of the Family Health Strategy. 300 subjects participated. Variable associations were significant (p<0.05) between religions with positive beliefs and attitudes in the neurological dimension (β: 1.040; p: 0.044); an increase in educational level had a significant relationship with negative beliefs and attitudes in the environmental/psychophysical dimension (β: −0.723; p: 0.040); having a child was significant for negative beliefs and attitudes in the environmental/psychophysical dimension (β: 1.120; p: 0.043) and for positive beliefs in the metaphysical dimension (β: −0,244; p: 0.028). This research contributed to the identification of factors influencing beliefs and attitudes towards childhood epilepsy existing in the studied Brazilian cultural context with the aim of contributing to a better implementation of actions directed to education in epilepsy.

## 1. Introduction

Epilepsy is the most common chronic neurological disease in childhood, affecting roughly 5 to 10 children in every 1,000, with profound repercussions in the cognitive, psychological and social areas [1,2].

Such impacts arise from interactions between multiple factors involving the clinical aspects of the disease and the adverse effects of drug treatments, as well as the expressive negative psychosocial connotations based on inappropriate beliefs and attitudes, which strengthen the social stigma and expose the children affected by the disease to discriminatory and prejudiced attitudes, negatively affecting their quality of life [3,4,5].

The lack of information regarding the disease is believed to be an important contributing factor to the persistence of enormous social stigma, beliefs and inappropriate behavior [2,5,6,7]. However, other factors can also influence the propagation of such actions, thus efforts aimed at identifying these are very important since these factors need to be considered when implementing educational actions to reduce the stigma, beliefs and negative attitudes of the general community towards the affected children.

In this context, the Family Health Strategy (FHS), for being set up as the gateway to health services and having greater proximity to users, stands out as a conducive environment for the implementation of actions directed towards epilepsy education, where it is essential that health professionals associated with such actions are aware of the factors capable of influencing the beliefs and attitudes of the population regarding the disease.

Thus, this study aimed to analyze the possible factors influencing the beliefs and attitudes of the general population towards childhood epilepsy among users of FHS units in the city of Crato, Ceará, northeastern Brazil.

## 2. Material and methods

This is a study with a quantitative approach, where the research was conducted in three units from the FHS located in the urban area of the municipality of Crato, in the region of Cariri, Ceará, Brazil.

The participants involved in the research were registered users at the FHS units, who were present and available at the time of data collection at the institution. In addition, the following aspects were also considered as inclusion criteria in the study:be aged between 18 and 59 years old and be literate(a).

The exclusion criteria were: people with allopsychic and autopsychic disorientation; those suffering from psychiatric disorders whichmay alter their understanding of reality; as well as sedative users which may experience changes to a greater or lesser extent in their motor or mental functions.

Participants were contacted in the FHS waiting room, where those who agreed to participate in the research and met the inclusion criteria were selected. The sample calculation was based on structural equation modeling techniques. The calculation is based on the maximum permitted levels for type I and type II errors,the acceptable limits for the root mean square error of approximation (RMSEA) (between 0.5 and 0.8) and the degrees of freedom of the model [8].

Data collection took place between September and October 2018. To this end, participants individually answered a questionnaire for socioeconomic characterization. Thereafter, participants were instructed to answer part II of the Brazilian version of The Epilepsy Beliefs and Attitudes Scale (EBAS) - Adult Version, in order to investigate their beliefs and attitudes towards childhood epilepsy [9].

The instrument is divided into: Part I with six questions addressing the participant’s general knowledge and experience with epilepsy; Part II containing a story highlighting the symptoms and behaviors during and after an epileptic seizure in a child, followed by 46 beliefs and attitudes associated with childhood epilepsy [9].

In part II, participants were instructed to read the story and select from a four-point Likert scale which of the following response alternatives represented the intensity/degree of their belief for each of the 46 items: (1) I don’t believe it, (2) I believe it a little, (3) I believe it a lot, (4) I completely believe it [9].

The 46 items are divided into three dimensions: neurological, metaphysical and environmental/psychophysical with 13, 7, and 26 items, respectively [5]. Additionally, the 46 items address both positive and negative beliefs and attitudes towards epilepsy [2].

Data obtained from the socioeconomic questionnaire were analyzed using descriptive statistics (absolute and relative values) with the JASP software (Version 9.0.1).

For the data obtained from part II of the EBAS-Adult Version instrument, a structural equation analysis [10] was performed with the Mplus software and divided into two steps. Firstly, a confirmatory factor analysis of the 46 items present in part II of the instrument was performed in order to validate the fit of this theoretical measurement model. For this, the quality of the fit of the 46 items with their neurological, metaphysical and environmental/psychophysical dimensions were evaluated. Secondly, an estimation of the structural model was performed, where this analyzed the relationship of the 46 items grouped in the three dimensions (neurological, metaphysical and environmental/psychophysical) with the variables: sex, religion, education, having a relative with epilepsy and having a child son. Variable having a child, we considered all participants who reported being a parent. For the having a child covariate, all participants who reported being a parent were considered.

In both structural equation models, theoretical and structural, the correlation coefficients of all paired combinations of the indicator variables were estimated according to the nature of the variable. The Weighted Least Squares estimation method [11] was used in all models since it is the most appropriate when using categorical variables, where the standardized direct and total effects were estimated [12]. The Comparative Fit Index (CFI) which indicates sample quality, the Tucker Lewis Index (TLI) which addresses the items’ number ratio for the sample number and the RMSEA which shows the residual value, were used to assess the fit of the model. An approximate value of 0.80 was considered to infer the fit of the model for the CFI and TLI [13], while RMSEA values from 0 to 0.08 were considered an acceptable fit [14].

In this research, the requirements of the Guidelines and Norms of the Research Involving Human Beings, regulated by Resolution 466/12 of the National Health Council, were met [15]. The project received an opinion of approval from the Research Ethics Committee, under number 2.895.570.

## 3. Results

### 3.1. Profile of the Survey Participants

As shown in (Table 1), 300 subjects aged between 18 and 59 years old, most of which were female (82.7%), participated in the research.

**Table 1.** Socioeconomic profile of the participants. Crato - CE, 2018.

| **Municipality** | **Institution** | **N** | **%** |
| --- | --- | --- | --- |
| **Crato – CE** | ESF | 300 | 100 |
| **Gender** |  |  |  |
| Female |  | 248 | 82,7 |
| Male |  | 52 | 17,3 |
| **Age Group** |  |  |  |
| 18-25 |  | 81 | 27,0 |
| 26-35 |  | 89 | 29,7 |
| 36-49 |  | 88 | 29,3 |
| 50-59 |  | 42 | 14,0 |
| **Marital Status** |  |  |  |
| Single |  | 155 | 51,7 |
| Married |  | 106 | 35,3 |
| Stable Union |  | 26 | 8,7 |
| Separate |  | 6 | 2,0 |
| Widow/Widower |  | 7 | 2,3 |
| **Non-Educated** |  |  |  |
| Incomplete Elementary School |  | 50 | 16,7 |
| Complete Elementary School |  | 16 | 5,3 |
| Incomplete High School |  | 39 | 13,0 |
| Complete High School |  | 128 | 42,7 |
| Incomplete Higher Education |  | 30 | 10,0 |
| Complete Higher Education |  | 37 | 12,3 |
| **Religion** |  |  |  |
| Evangelical |  | 54 | 18,0 |
| Catholic |  | 208 | 69,3 |
| Spiritualism |  | 1 | 0,3 |
| Umbanda |  | 2 | 0,7 |
| Jehovah's Witness |  | 1 | 0,3 |
| No religion |  | 34 | 11,3 |
| **Occupation** |  |  |  |
| From the Home |  | 86 | 28,7 |
| Student |  | 37 | 12,3 |
| Teacher |  | 18 | 6,0 |
| Other professions |  | 150 | 50% |
| Unemployed |  | 9 | 3,0 |
| **Monthly family income** |  |  |  |
| < 1 minimum wage |  | 198 | 66,0 |
| 1 - 2 minimum wages |  | 64 | 21,3 |
| 2 - 3 minimum wages |  | 30 | 10,0 |
| > 3 minimum wages |  | 8 | 2,7 |

Source: Search Data, 2018.

Regarding the marital status of the participants, 51.7% reported being single. Most participants (42.7%) had completed high school, 69.3% claimed to be of the Catholic religion and 66.0% received up to 1 minimum wage in monthly income. Fifty-seven professions were cited with most participants, corresponding to 28.7%, performing household activities.

### 3.2. Validation of the Measurement Model

The general fit indices of the model presented excellent values (CFI=0.90; TLI=0.91; RMSEA 0.05) indicating data adherence to the theoretical construct evaluated by the EBAS - Adult Version instrument.

Figure 1 shows the factor weightings where most items were acceptable, ranging from 0.32 to 0.78with some exceptions. In the neurological dimension, the factor weightings ranged from 0.32 to 0.54, with the exception of items 15, 35 and 37 with low factor weightings, obtaining 0.10, 0.17 and 0.29, respectively. In the metaphysical dimension, the factor weightings ranged from 0.53 to 0.68, except for item 32 with 0.03. In the environmental/psychophysical dimension the factor weightings ranged from 0.33 to 0.78, with the exception of items 1, 17, 22 and 36, with low factor weightings, which obtained 0.23, 0.05, 0.04 and 0.20, respectively.

**Figure 1.**
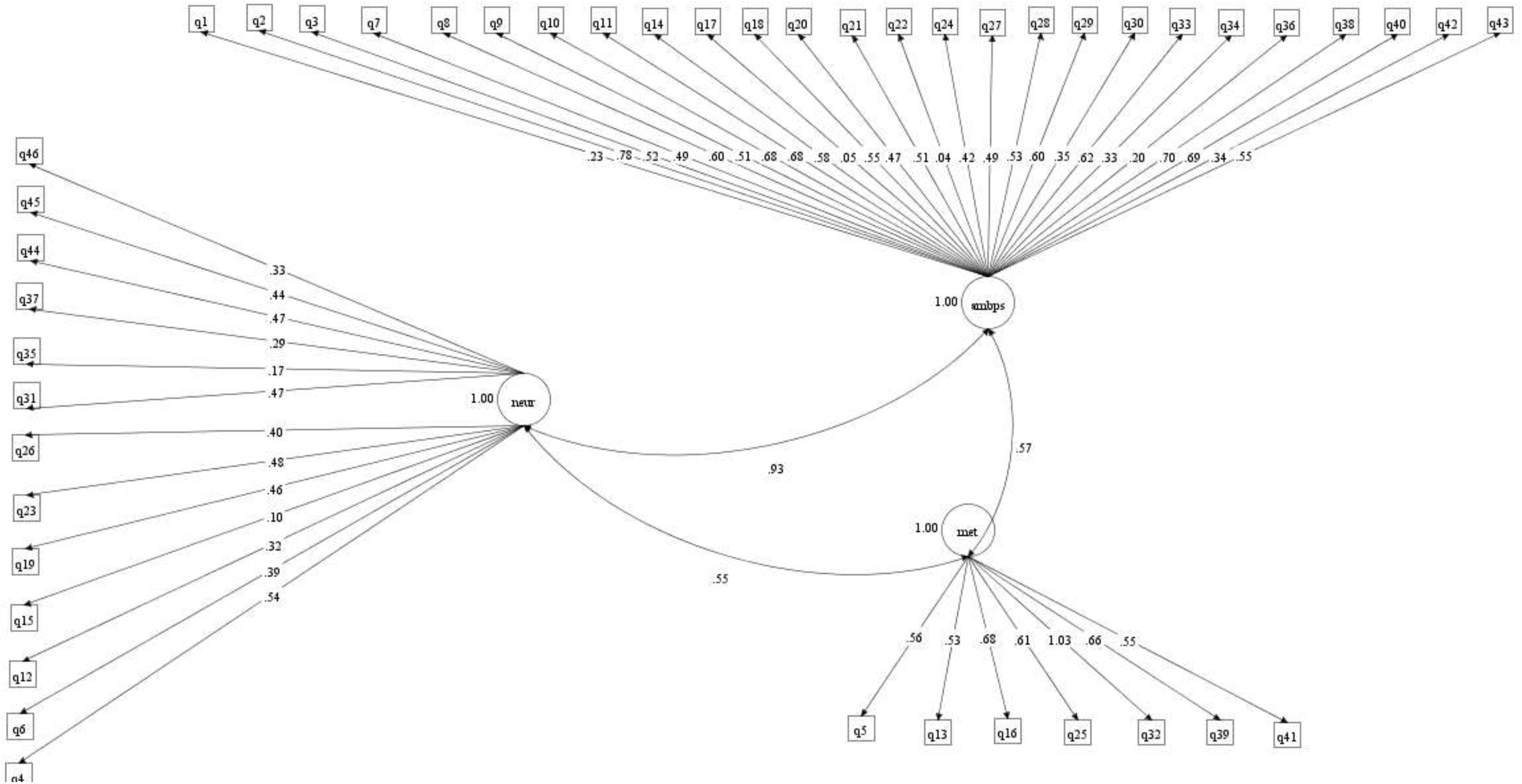
Measurement model from part II of the Brazilian version of the EBAS - Adult Version. Caption: q: question; amb/pis: environmental/psychophysical; met: metaphysical; neu: neurological. Source: Search Data, 2018.

Therefore, as most items obtained acceptable factor weightings, presenting a good adjustment for their dimensions and the overall adjustment indices were also excellent, items which presented unacceptable results were not to excluded from the scale as they were considered valid from a theoretical viewpoint. Thus, no changes were made to the instrument, keeping the causal structural equation analysis of all EBAS - Adult Version items in Portuguese the same.

### 3.3. Structural Model: Validation and Association between Variables

In the structural model validation, the analysis indicated that the general adjustment indices CFI/TLI/RMSEA were acceptable presenting values of 0.90, 0.91, 0.04, respectively, thus ensuring the reliability of this model, which includes the association of the 46 items distributed across the three dimensions (Neurological, Metaphysical and Environmental/ Psychophysical) from Part II of the instrument with the variables sex, religion, education, having a relative with epilepsy and having a child (figure 2).

**Figure 2.**
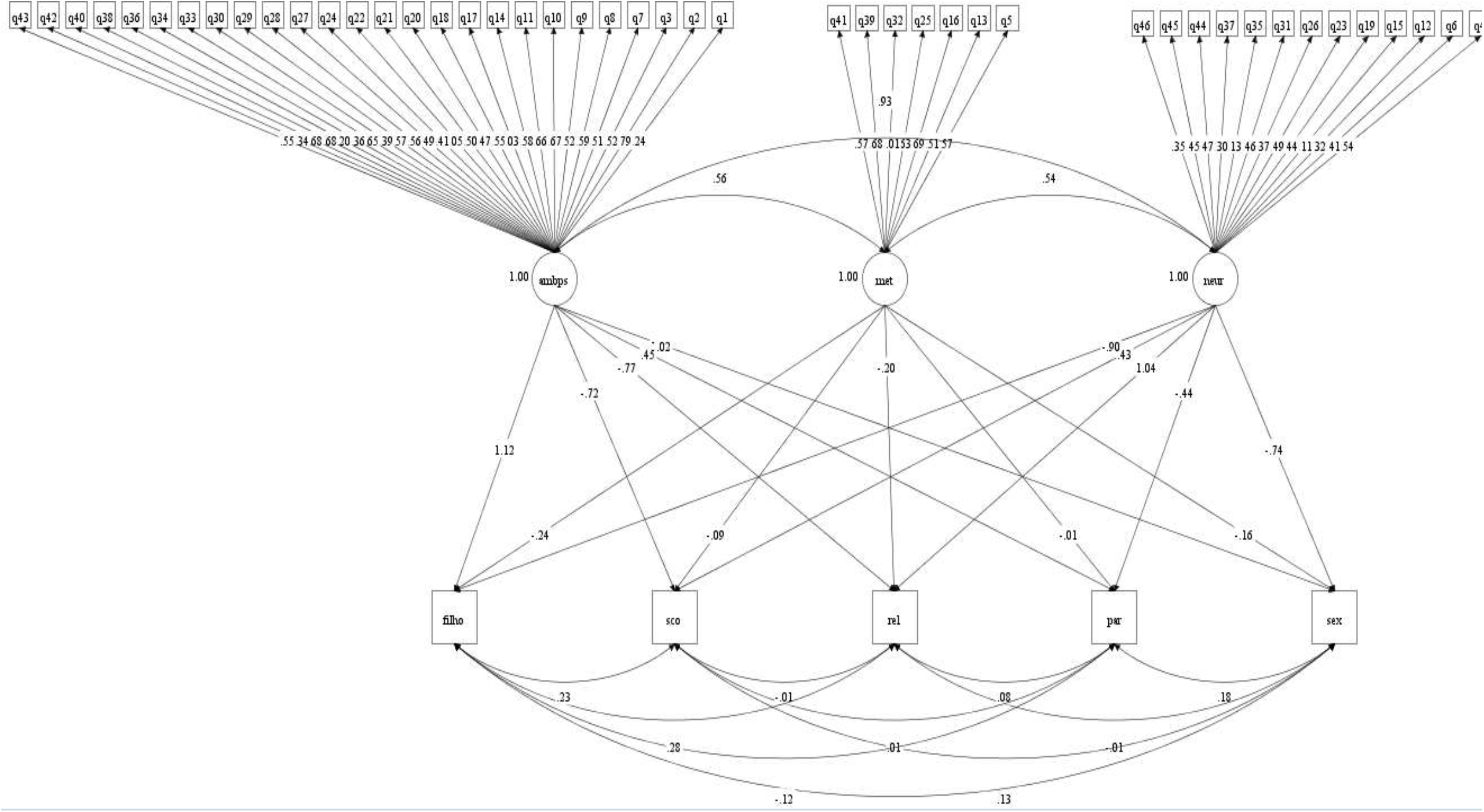
Structural model from part II of the Brazilian version of the EBAS - Adult Version. Caption: q: question; amb/pis: environmental/psychophysical; met: metaphysical; neu: neurological; filho: having a child son; sco: education; rel: religion; par: having a relative with epilepsy and; sex: gender. Source: Search Data, 2018.

Figure 2 shows the association results were significant (p<0.05) for three variables (religion, education and having a child) in different dimensions (neurological, metaphysical and environmental/psychophysical). The variables sex and relatives did not present values considered significant.

The relationship between religion and the neurological dimension was significant and positive (β: 1,040; p: 0.044). As for the level of education, a significant and negative relationship with the environmental/psychophysical dimension (β: −0.723; p: 0.040) was observed, whereas for the having a child variable, a significant and positive relationship with the environmental/psychophysical dimension (β: 1.120; p: 0.043) was observed, as well as a significant and negative relationship with the metaphysical dimension (β: −0.244; p: 0.028).

It is noteworthy that only one participant was the mother of a child with epilepsy, the other participants reported their child did not have the disease.

## 4. Discussion

The results obtained in this study showed that having a religion increased the relationship between beliefs and positive attitudes pertaining to the neurological dimension of the EBAS-Adult version instrument, which are supported by a scientific basis when dealing with the neurological and genetic genesis of epilepsy [16].

It is noteworthy that 88.6% of participants declared having a religion, with the majority being Catholics (69.3%). In the neurological dimension, positive beliefs and attitudes surrounding epilepsy were due to a belief that epilepsy can be caused by brain abnormality, brain damage at birth and genetic defects, as well as having the physician as the best professional to care for the affected child [5].

Therefore, such a significant influence of religion on increasing beliefs surrounding epilepsy as a disorder of the central nervous system may be inferred as a remarkable change in the religious view of the causes of epilepsy, these being previously associated with supernatural influences (demonic possession) and higher powers (God’s will) to a more centered view on the neurological basis of epilepsy being caused by changes in the central nervous system [17].

Corroborating with these findings, Rafael et al. [18] found the attitudes of the general population showed that most participants agreed that epilepsy is not a supernatural disease, correctly identifying brain injury as one of its possible causes, in addition to not considering it to be contagious.

As for the influence of higher educational levels with an increased association with negative beliefs and attitudes in the environmental/psychophysical dimension, the interference of this factor withan increase in non-medical (popular) beliefs and attitudes associated with the influence of changes in time in the occurrence of epileptic seizures (eg sudden changes in weather or lunar phases) and those associated with psychophysical aspects (eg possession by evil spirits and having seizures when angry) was evidenced [16].

However, it is noteworthy that the largest representativeness from the sample in this study were from people who did not have a higher educational level (77.7%), and because these participants did not have a large amount of instruction compared to those who concluded higher education, this may be an important aspect strengthening their negative beliefs towards childhood epilepsy. However, when analyzing studies published at national and international levels, no consensus in the literature has yet been observed regarding the influence of educational levels on beliefs and attitudes towards epilepsy, thus showing the lack of information regarding the disease affects all educational levels and may be a factor of considerable influence on inappropriate knowledge. Some research shows that inadequate knowledge is more closely associated with lower levels of education, while other studies conducted with people with higher education also show the remarkable presence of inappropriate beliefs [2,5,18,19,20,21].

In this study, being the parent of a child influenced increased negative beliefs and attitudes with the environmental/psychophysical dimension, such as believing a child may have epileptic seizures due to: possessions by evil spirits, playing in the sun for a prolonged time, sudden changes in weather (becoming very hot/cold/humid/rainy), certain foods/drinks, changes in lunar phases or traveling in vehicles without air circulation [5]. While also influencing positive beliefs in the metaphysical dimension such as believing that faith in a higher power helps coping with epilepsy [5].

This finding may be associated with the affective component of beliefs, since according to Wright et al. [22], beliefs comprise three components: cognitive, affective and behavioral; with the affective component being responsible for instigating the occurrence of favorable or unfavorable feelings towards an object, event or behavior.

Thus, from the results of this study, being the mother or father of a child possibly favored empathy towards children with epilepsy, and it is noteworthy that playing this role may involve a feeling of overprotection that leads to an attachment to a favorable belief or faith in a higher being,given an individual’s impotence in coping with a chronic disease such as epilepsy, however which can also strengthen their negative beliefs towards the environmental and psychophysical aspects of the disease, as a way of trying various ways to protect the child from further harm to their health.

Similar results were obtained in the study conducted by Zanni [5] when applying the Brazilian version of the EBAS - Adult version to 56 parents of children with epilepsy, where the results demonstrated adequate beliefs in the Metaphysical dimension (Averages: 2.7 and 3.0) in item 41 which addresses faith in a higher power to help cope with epilepsy. While in the Environmental/Psychophysical dimension (Averages: 2.7 and 2.9), inappropriate beliefs and attitudes surrounding children having epileptic seizures when playing in the sun for a prolonged time, due to sudden changes in the weather, food/drinks or changes in lunar phases were observed.

Thus, considering the influence of the Brazilian cultural context, specifically in the Cariri region, on the factors investigated herein and which demonstrated to interfere with the knowledge of the target population is emphasized.

Gajjar [16], conducted a study focusing on the beliefs and attitudes towards epilepsy in children, which indicated the cultural characteristics of a group, personal experiences with epilepsy, exposure to acculturation and/or interactions with different cultures were found to be fundamental factors in the construction of different beliefs surrounding childhood epilepsy. Bartolini, Bell and Sander [23] also devoted themselves to reviewing multicultural challenges in epilepsy, with the aim of raising awareness to the importance of socio cultural knowledge.

The knowledge surrounding childhood epilepsy may be influenced by different factors and it is, therefore, appropriate that these elements receive attention from healthcare professionals, who in turn, should consider these during the implementation of actions focused on healthcare education surrounding epilepsy.

## 5. Conclusion

The findings from this study emphasize that having a religion, a higher educational level and being the parent of a child, were shown to be factors influencing the beliefs and attitudes of the target population regarding childhood epilepsy.

It is noteworthy the present research was limited to approaching users of FHS units only in an urban area. However, given the scope of the addressed theme, further research is needed to encompass individuals from different contexts, identifying other variables capable of influencing the dissemination of negative beliefs and attitudes towards the disease and how healthcare professionals may develop and disseminate successful healthcare education al strategies within healthcare education for epilepsy care.

## Conflicts of interest

The authors declare that there are no conflicts of interest.

## Acknowledgments

The contributions of Luanna G. Silva, data collection and article editing; Paulo F. R. Bandeira, by analyzing the research data; Cicero D. Alencar, aid in data collection; Gyllyandeson A. Delmondes, Simone S. Damasceno e Marta R. Kerntopf, general evaluation and correction of the article; Izabel C. S. L. Beltrão, final review of the article.

